# Contractile Vacuole and Papilla drive Cyst/Telotroch transition in *Vorticella microstoma*

**DOI:** 10.1101/2025.03.06.641897

**Authors:** Eric Galiana, Marie-Line Kuhn, Agnès Attard

**Author notes:** Corresponding author.* Eric Galiana. Institut Sophia Agrobiotech, 400 route des chappes, BP 167, 06903 Sophia Antipolis Cedex FRANCE.

## Abstract

The cyst of the protist ciliate Vorticella efficiently differentiates into teletroch, the free-swimming stage. Here, using video microscopy and quantitative imaging we establish that the cyst differentiation follows two strict temporal and spatial patterns. The temporal pattern is initially marked by the functional formation of the contractile vacuole, which then discharges its fluid into the neoformed cytopharynx and, in a third stage, into a membrane invagination which, to the rhythm of the vacuole, differentiates into an oral cavity exhibiting a polarized cilia array revealed by α-tubulin immunostaining. Two poles delineate the spatial pattern. The apical pole is defined by the position of the pre-existing papilla, which determines the site of oral cavity formation directly below it. The second basal pole is placed at the rupture point of the cyst wall. It is characterized by β-actin accumulation and allowed release of the telotroph according to its basal-apical polarity. Both the temporal and spatial patterns are impaired by concanamycin A treatment, a specific inhibitor of vacuolar type H^+^-ATPases altering functions of contractile vacuoles. The findings indicate that cyst to telotroch transition is predetermined by the location of the papilla on the cyst wall and under the control of a functional contractile vacuole. We propose that *Vorticella* cyst is a simple single-cell model for investigating basic principles of integrative neogenesis of organelles in Protozoa, and of apical-basal cell polarity in Eukaryotes.

## INTRODUCTION

Van Leeuwenhoek (1677) observed with his light-refracting lens microscope the first microbes, presumably a *Vorticella*. He described them as poor little creatures living in rainwater, having a round or oval body, moving or harpooning themselves towards the extremity, where they possess a contractile tail (stalk), and clumping together in a few filaments. The *Vorticella* genus is now a gathering of more than 100 different sessile peritrich ciliates (Sun et al, 2013). They evolve in marine, freshwater and terrestrial environments, moving or being attached to various substrates such as plant/animal surfaces and appendages, biofilms, flooded objects (Liao *et al*., 2018; Galiana *et al*., 2011; Davis and Gloege, 2024). They have been characterized as dominant protozoan species putatively playing a role in fertilization on N supply in rhizosphere soil during the initial growth of *Beta vulgaris L.* (Zeng *et al*., 2020), and as potential biocontrol agents against plant-parasitic nematodes (Rostami *et al*., 2022).

At the cellular level, they develop at least in three single-cell forms: free-swimming telotrochs, sessile stalked trophonts and cysts, the last ones being formed under adverse environmental conditions such as lack of nutrients or oxygen, drought, osmotic stress and extreme temperatures (Finley and Lewis, 1960; Bramucci and Nagarajan, 2004; Ryu *et al*., 2017). The free-swimming telotroch (0.2-1 mm/s in speed) has generally an inverted bell-shaped morphology with an apical part surrounded by at least two rings of cilia emerging from the peristome, which surrounds the open oral cavity/mouth. Immediately behind, the cytopharynx, resulting from a plasma membrane invagination, connects together the oral cavity, particularly, to the food vacuoles, the digestive organelles, as well as the single contractile vacuole (CV). The osmoregulatory CV presumably emptying its contents into the cell’s cytopharyngeal region through an elongated tube (Allen and Naitoh, 2002). The cell show the presence of a micronucleus and a polylobar macronucleus and the other eukaryotic organelles. The trophont sessile form has the same morphology and exhibits a retractile stalk (100-150 µm) with which the cell is attached to a surface. The water vortex flow generated by oral cilia beating results in drawing bacteria food particles toward the peristome and then to food vacuoles ingesting particles using phagocytosis. *Vorticella* species with their trophont-stalk contracting almost instantly (10-20 mm/s) and the high swimming speed (0.2-1 mm/s) of the free telotroph are among the fastest cellular micromachines making them highly attractive models for studying movements from hydrodynamic, cellular and molecular point of views (Kamiguri et al, 2012; Ryu *et al*., 2017; Lisicki et al, 2019; Lupatelli *et al*., 2023).

Perhaps because cyst formation is a slower process, the dynamics of encystment occurring upon adverse conditions, and excystment, allowing telotroch release from the cyst, are less documented. (Warren, 1986). Cyst formation results to the formation of an envelope, the cyst wall, encapsulating a diffuse cytoplasm/hyaloplasm with indistinct cilia, peristome, cytopharynx and vacuoles. Upon favourable conditions, excystment is induced and characterized by cyst differentiation, functional cilia and CV, with osmolarity as a main regulatory factor (Finley and Lewis, 1960).

In protists, osmoregulation is achieved by CV which maintains osmotic and ion homeostasis (Plattner, 2013, 2015; Jimenez *et al*., 2022; More *et al*., 2024). The cycle of CVs involves three sequential events: diastole (slow extension of the vacuole gradually filled with excess water drained into the endoplasm); systole (rapid vacuolar contraction); and expulsion (the release of vacuolar fluid from the CV outside the cell). To maintain osmotic and ion homeostasis through extrusion of water and excess of ions, the CV may exploit the H^+^-gradient generated by the organelle-resident Vacuolar ATPases (V-type H^+^-ATPase) (Patterson, 1980; Plattner, 2013, 2015). The V-ATPase is highly conserved in eukaryotic organisms and is assembled into two structural multimeric protein complexes: a transmembrane V0-proton pump and a cytoplasmic V1-ATPase that can be reversibly detached and reattached (Wassmer *et al*., 2006; Wang *et al*., 2020). In protists as *Dictyostelium* and *Phytophthora* (Temesvari et al, 1996; Mitchell and Hardham, 1999) the function of vacuolar contraction is impaired by concanamycin A, a specific inhibitor of V-ATPases, binding to the V0 subunit c (Huss et al, 2002).

To date and for a long time ago, knowledge of the *Vorticella* cyst still remains meagre (Warren, 1986). The underlying machinery controlling encystment and excystment remains unexplored since the first study of Finley and Lewis (1960). Here, we combine quantitative imaging with drug-inhibited V-ATPase activity to characterize the dynamics of excystment and telotroch release from the *Vorticella microstoma* cyst induced in response to hypo-osmotic conditions. We established that before release, oral cavity and cilia neogenesis, as well as cyst wall deformation follow the redifferenciation of the CV and that their formation is tightly synchronised both spatially and temporally with the acceleration of the pulsatile rhythm of the diastole-systole-expulsion sequence. We show that inhibition of V-ATPase-mediated proton pumping causes drastic deficits in telotroch release, speed motion as well as cilia organisation and distribution. We propose that *Vorticella* cyst is a simple model for investigating basic principles of integrative neogenesis of organelles in Protozoa, and of basal-apical cell organisation in eukaryotes.

## RESULTS

### Excystment main features

Encystment and excystment have been reported in several ciliated protists (Li et al, 2022; Jin *et al*., 2024), in few species within the *Vorticella* genus (Warren, 1986) among which *V. microstoma* (Finley and Lewis, 1960) that was investigated in this work. Encystment was induced at a rate of 100% after incubation of *V. microstoma* cells in V8 medium in long-term-culture (10–14 days). At these time-points of incubation, no trophont nor telotroph could be detected with binoculars. The dynamics of encystment was not further examined in this study. To further study the dynamic of excystment, we applied a hypo-osmotic shock to this cyst population. In these conditions, the rate of excystment was estimated at 30% to 50% within 1-2 hours. At this time, microscopic observations allowed to establish that the cell released were telotrophs either swimming freely or not. Further investigation on excystment dynamics focused on telotrophs that were not yet swimming and were almost exclusively localized immediately adjacent to the empty cyst wall from which each had emerged. In the results that follow, the timing assigned to each event described refers to the moment (t=0) when osmotic stress was applied.

Just before the stress, all *Vorticella* material consisted of perfectly circular and encapsulated cysts (⌀28.8±0.9 µm) with a papilla located at one of its poles as illustrated in Fig. 1A and 1B. At this time (t=0), an immunostaining for the α-tubulin subunit revealed an intracellular and diffuse staining homogeneously distributed in the cytoplasm, suggesting that microtubules were not or poorly polymerised and cilia resorbed (Fig. 1B). Around t=30 min, re-differentiation was initiated as attested by the formation of the contractive vacuole (Fig. 1C) and the cytoplasmic staining for α-tubulin which became punctuated indicating αβ-tubulin oligomerisation in clusters (Fig. 1D). From around t=30-60 min, telotroch release from cysts took place in number. The turgor pressure generated by the telotroph size increase stretched the cyst wall to its maximum capacity up to a breaking point. Cells exhibited the characteristic telotroch bell morphology decorated by cilia (Fig. 1E and 1G). As previously described, their size (1595,5 ± 58,8 µm^2^) was significantly increased (p=0,0005; n=14) by 2,4 times compared to cysts (571,2±36,9 µm^2^).

**Figure 1.**
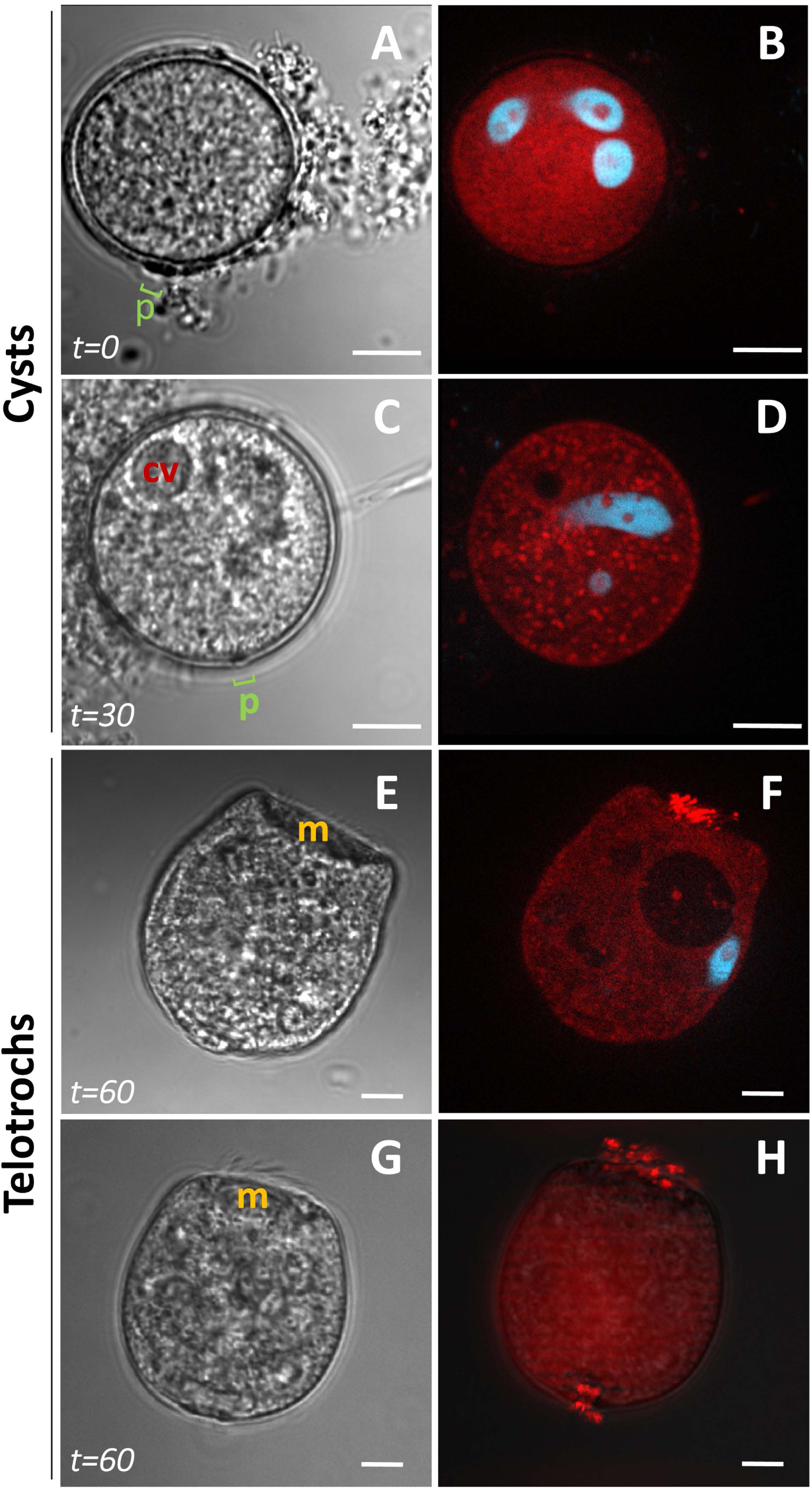
Cysts and telotroch Morphologies. (**A, C, E, G**) Representative images of phase-contrast micrographs 0, 30 and 60 min after induction of the hypo-osmotic stress: cysts (**A, E**), telotrophs (**E, G**). (**B, D, F, H**) Representative images of immunofluorescence of α-tubulin (red) and DAPI staining of polylobar nuclei (blue) in cysts (**B, D**) and telotrochs (**F, H**). Scale bars: 10 µm.

Immunostainning for α-tubulin (Fig. 1F and 1H) clearly showed occurrence of microtubule polymerisation and cilia neogenesis at the apical side (the oral mouth) of the telelotroph (Fig. 1E and 1G), and at its basal side (Fig. 1G). The rupture point of cyst wall at release was detected at the opposite side of the papilla when this one was visible in the plan (Supplementary Fig. 1). The position of this point was inscribed in the half-circle opposite to the location of the papilla. This indicated that at the time of rupture, the pressure is exerted differently along a particular cellular axis. Another interesting point is congruent with this hypothesis: systematically, the release of the cell took place according to its apical-basal polarity, its basal end engaging first (Supplementary Fig. 1).

### Tracking early organelles neogenesis

To examine more precisely, the spatio-temporal dynamics of organelles neogenesis (contactile vacuole, oral mouth, cytopharynx, cilia, cyst wall deformation) within the cyst, samples were positioned in the chamber just after the hypo-osmotic stress. Cysts and excystment were captured at 5 frames per second until telotroch release (Fig. 2; movies 1, 2 and 3). The time series images were processed by kymography. Figure 2 summarises the occurrence of sequential events observed.

**Figure 2.**
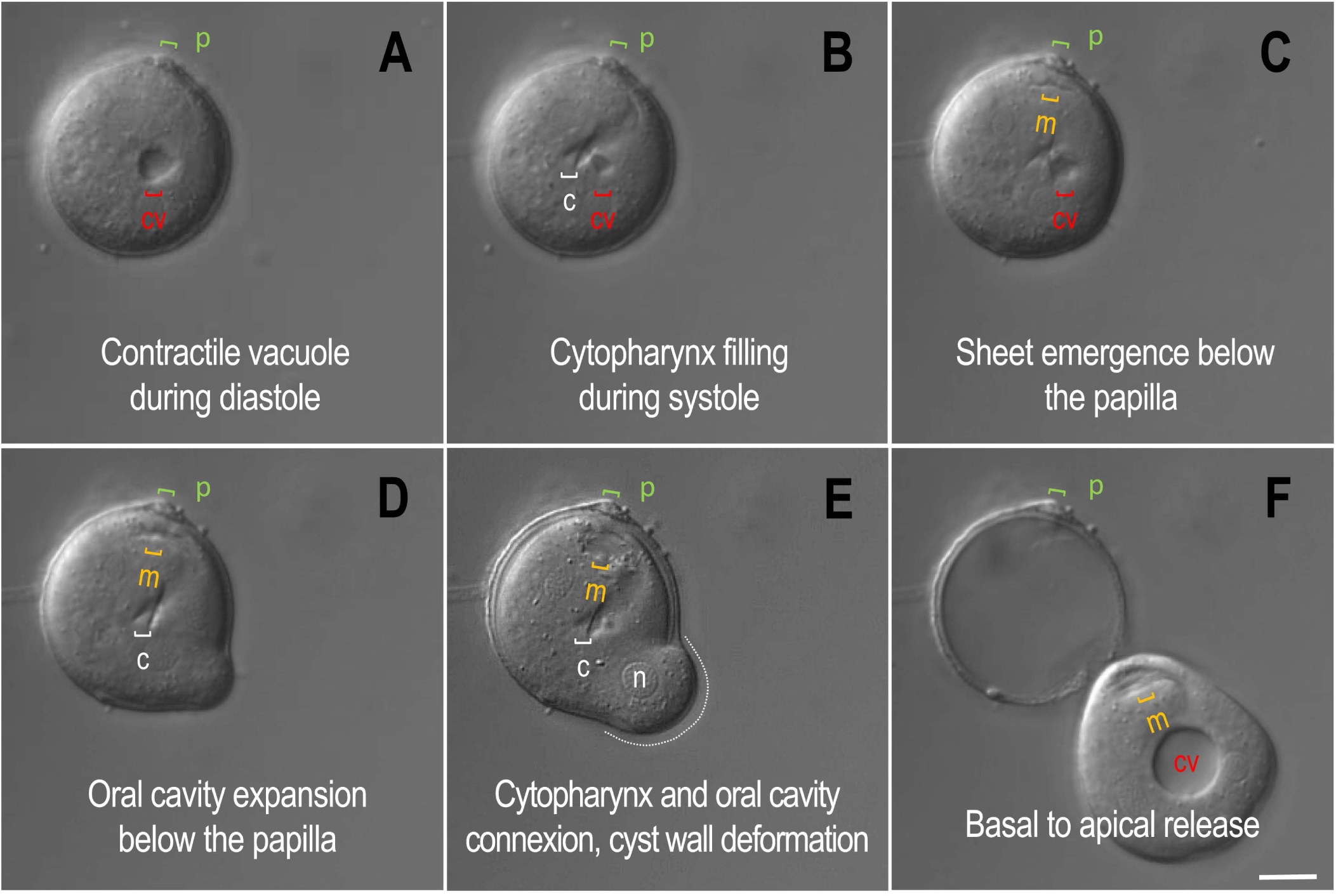
Time-lapse imaging of excystment. (**A-F**) A representative sequence of main events observed during cyst differentiation extracted from the Movie 1. In this movie t=0 was undetermined. Annotation indicates the localisation of the different structures : (p) papilla; (cv) CV; (c) cytophrynx; (m) oral mouth/oral cavity; (n) nucleus; the dotted white line identified the location of cyst wall deformation. The micrographs illustrate different interconnected steps of cyst differentiation: (**A**) the CV during diastole; (**B**) the cytopharynx filling during systole; (**C**) the emergence of a thin sheet beneath the papilla; (**D**) the oral cavity expansion from the thin sheet; (**E**) the cytopharynx and oral cavity connexion, and the cyst wall deformation; (**F**) Basal to apical release. Scale bar: 10 µm.

For the CV, we retrieved from kymograms the number of pulsations before release, kinetics of beating frequency, size of the vacuole at the end of systole (Fig 3). The data show that the time point of the first pulse was variable and detectable quite a long time [7-21 min range] after stress induction (Fig. 3A,B). The total number of pulse before release was constant (41±4, Fig. 3C). Within a range between 0,01 and 0,05 Hz, the contractile frequency distribution overtime reveals for all samples the same pattern: first slow pulsations with a progressive increase in pulse frequencies and then a decrease for the last 5 to 10 ones before release (Fig. 3D and Supplementary Fig. 2). These variations are mainly due to variability in diastole duration, while the duration of each systole appears to be very short and constant, estimated in these experiments at 0.8-1 s. As illustrated in Fig 3A, the CV increased in size with time reaching its maximal volume just before release. The overall size was not quantified with kymographic data because vacuole position related to the selected line along the diameter could change marginally due to low flow or Brownian motion acting on the cyst. Changes in vacuole diameter size were manually quantified at three steps: at the first (C**_f_**) and last (C**_l_**) pulse detected within cyst and immediately after release within telotroph (T) (Fig. 3E). At C_l_, diameter was approximately 1,9 (±0,5)-fold larger than at C_f_, and 1,6 (±0,4)-fold larger at T than at C_l_. Thus, we can approximate that during differentiation from cyst to telotroch the volume of the CV increased 15 to 20-fold time.

**Figure 3.**
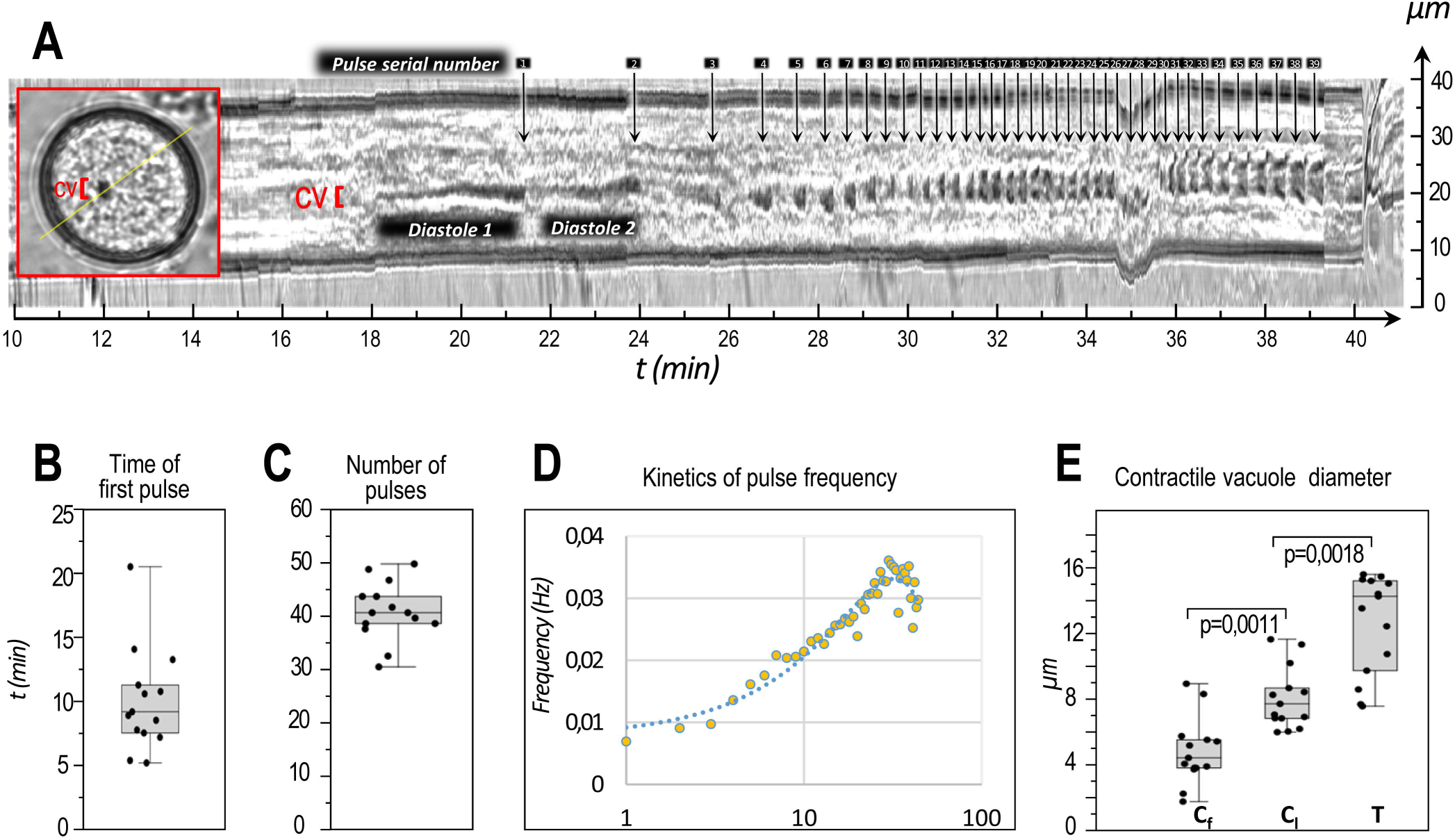
Kymogram of differentiation from cyst to telotroph. (**A**) Kymograph generated from Movie 2 showing the kinetics of CV pulses. Inlet shows the position of the section-line selected and the localisation of CV during the first diastole detected. On the kymogram are indicated : (i) the position of CV, (ii) the duration of the first and second diastole detected and (iii) by vertical arrows each pulse serial number positioned arbitrarily at the end of each diastole indicated (**B**) Boxplot of time for detection of a first pulse (n=14). (**C**) Boxplot of number of pulses counted from t=0 to telotroch release (n=14). (**D**) Mean of vacuole contraction frequencies at each pulse time series number (n=9). (**E**) Boxplot of vacuole diameter: within cyst at the first (C**_f_**) and last (C**_l_**) pulse detected and within telotroph (T) immediately after release (Fig. 3E, n=14).

Cytopharinx neogenesis was difficult to date precisely. However, when cytopharinx structures were first visualized in several successive time lapse sequences they clearly appeared only in correlation with the ability of vacuole to contraction (movie 1, 2 and 3). Moreover, part if not all the vacuole fluid discharge flew into the cytopharinx at systole. Indeed cytopharyngeal volume began to increase on the same frame on which for the first time at the end of diastole, systole started as attested by the decrease of the vacuole volume (Fig. 2A-B and Fig. 4). This increase reached its maximum in the same range of systole duration 0.8-1,2 s (Fig. 4A-D) even if here, the frequency of image acquisition did not allow a finer temporal resolution of these two rapid events.

**Figure 4.**
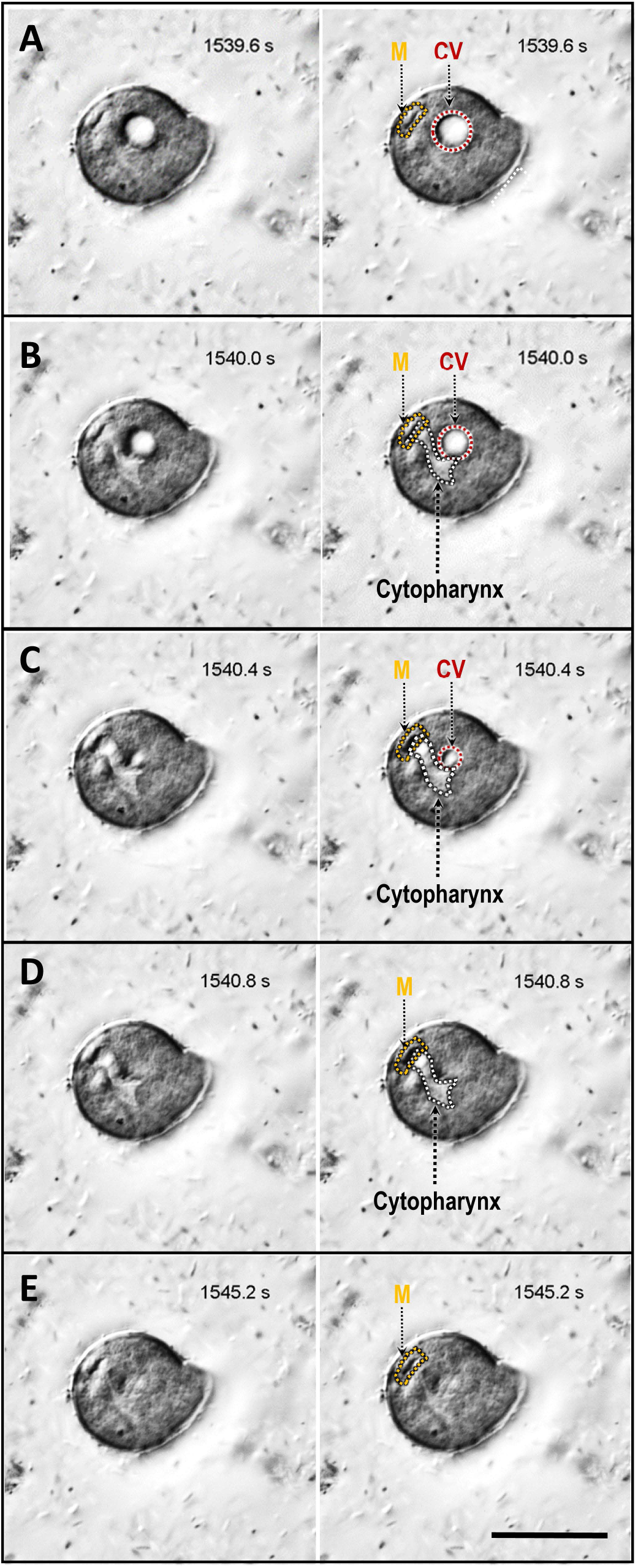
Time-lapse imaging of systole and cytopharynx filling. (**A-D**) Static images in duplicate, without (left) and with annotation (right), and taken in Movie 3 at 0.4-s intervals. Images show the proximal continuity between oral mouth (M), cytopharynx and CV (CV) localisation during the last systole before telotroch release. (**E**) Image shows the last image before a new diastole occurs. Scale bar, 30 μm.

Remarkably, our refined temporal analysis also revealed that oral cavity/mouth arise from a thin sheet, that could correspond to a plasma membrane invagination, whose expansion was cadenced by the vacuole contraction (Movie 1, 2 and 3). This primitive thin sheet is first visualized within the hyaloplasm directly and always beneath the papilla (Fig. 2C; Movie 1), probably involving a specialized cell cortex area for anchorage. At this stage, no cilia are visible and appear neither formed nor functional. The sheet gradually individualizes by rhythmic expansion, evolving into a cavity under successive repetitions of vacuole pulsations. The vacuole might empty its fluid into the cytopharyngeal region which joins the oral cavity by forming a tube (Movie 1). It is also noteworthy to note that when a kymogram was drawn based on a section line crossing organelles from the papilla to the vacuole (Fig. 5A, B), the oral cavity enlargement timely and perfectly coincided with the diastole/systole transition (Fig. 5C). The progressive enlargement could only be detected at the late stages of differentiation. At these stages, the emergence of cilia was detected as shown in Figure 2E-2F, and by the perceptible cilia vibration as the cavity expands just before and after release (Movie 1 and 3). Once formed, this cavity constitutes the oral mouth which appears to remain closed throughout the cyst differentiation-telotroch differentiation sequence.

**Figure 5.**
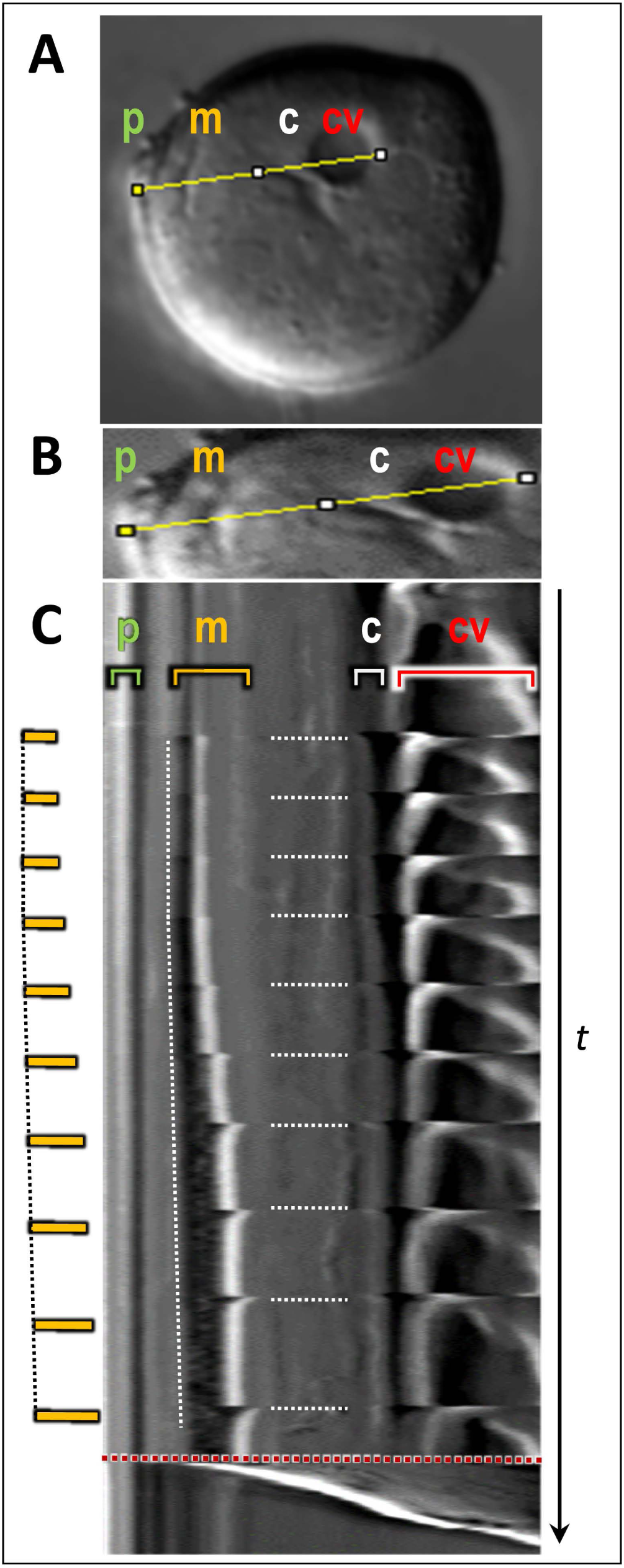
CV punctuates the tempo of enlargement of the mouth being formed. (**A, B**) images indicating how the section line was positioned to generate a kymogram. The section line crosses the papilla (p), oral cavity (m), cytopharynx (c) and CV (cv). (**C**) The image shows the final part of the kymogram including the last eleven contraction of the vacuole before release. Horizontal dotted white bars underline the perfect synchronisation between CV systole, filling of the cytopharynx and progressive and rhythmed oral cavity enlargement. Bold yellow bars illustrate the opening increment of the oral cavity, perfectly rhythmed in a phase pattern with the last vacuolar contractions occurring before release which is indicated by the red dotted line.

### Deformation and rupture of cyst wall

At latest stages of cyst differentiation, cyst wall deformation was initiated over a long and variable time [10-200s] before release (Fig. 6A). Sequence images analysis revealed that the cyst wall firstly underwent plastic deformation and then rupture. The plastic deformation exerted all along this time while the rupture was not quantified since it occurred in less than 0.2 s, the resolution time of the movies. Plastic deformation resulted from a protrusion at the leading edge taking place in an angular position located in the semi-circle opposite to the papilla coordinates (Fig. 6B) positioning the rupture point for the telotroph release which will follow (Supplementary Fig 1). The plastic deformation resulted with time in an increase in the perimeter of the cyst (Fig. 6C) that could reach just before release 24,2±7,9 µm (Fig. 6D) illustrating the significant elastic/elongation properties of polymers of the cyst wall. Deformation and rupture of cyst wall were associated with local β-actin accumulation revealed by immunostaining just before (Fig. 6 E-G) or during release (Fig. 6 H-J) suggesting that protrusion at the leading edge requires actin filament recruitment.

**Figure 6.**
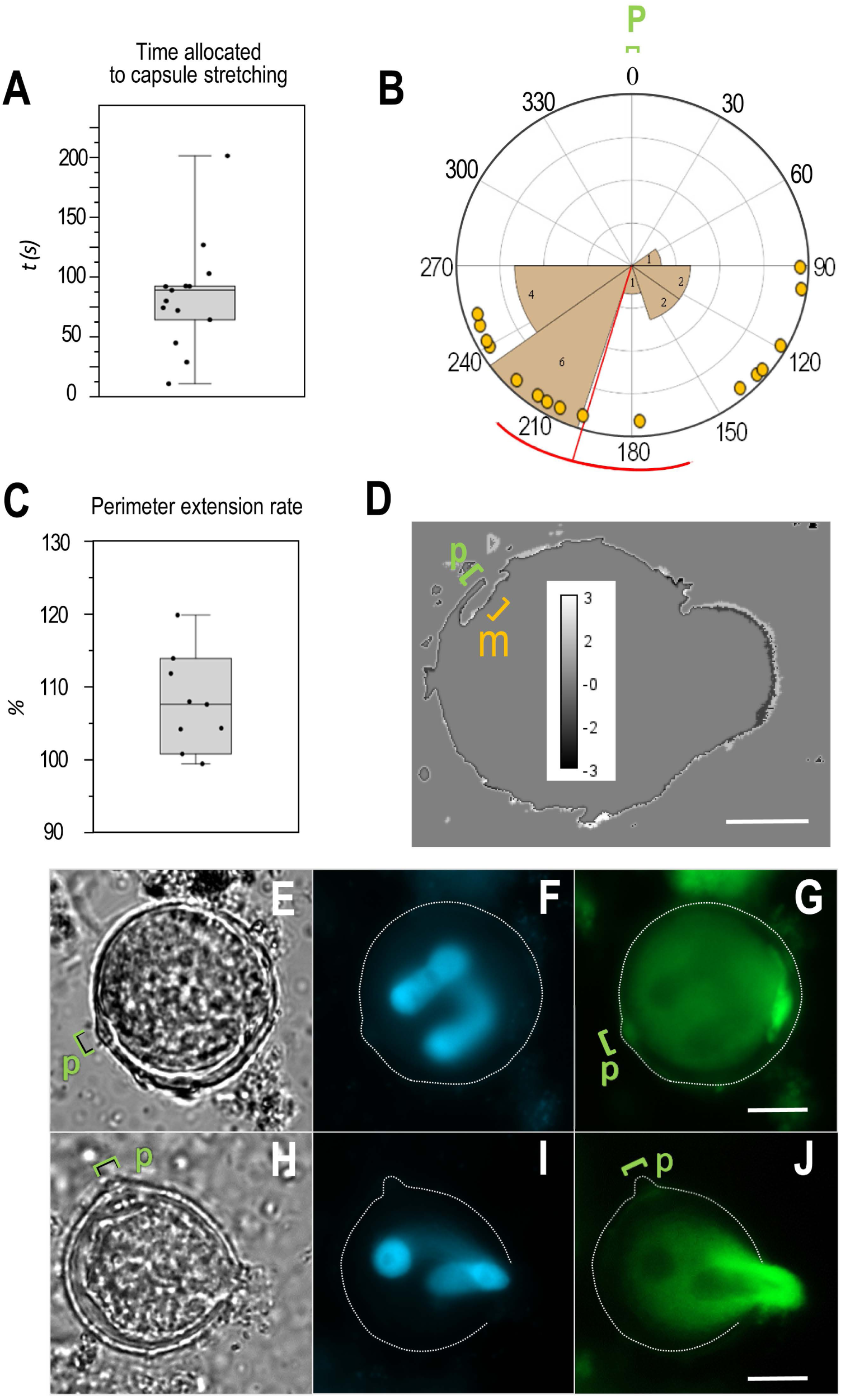
Deformation and rupture of cyst wall. (**A**) Time allocated to cyst wall stretching from protrusion emergence to release (n=14). (**B**) Polar plot of angle between papilla position and the rupture point. Range of angles between the half-line passing through the midpoint of the papilla (p; θ=0°) and the center of mass of the cyst, and the half-line passing through the center of mass and the midpoint defined at the site of protrusion emergence initiating the plastic deformation and then release. This range only contains obtuse angle values (n=16). (**C**) Percentage of increase in cyst size resulting from the ratio between the perimeters calculated at the time point just before release and before protrusion occurred (n=8). (**D**) Compute curvature of a perimeter cyst just before release showing local deformation of the cyst wall and the location of the papilla (p) and mouth (m). Calibration bar indicates value for point curvature (µm^-1^). (**E-J**) Phase contrast (**E-H**), DAPI staining of polylobar nuclei (**F, I**) and β-actin immunostaining (**G, J**) of cysts during release. Scale bars, 10 μm.

### Concanamycin A inhibits cyst differentiation and telotroch release

To initiate exploration of machinery controlling excystment, we assessed the function of the CV complex, the first organelle formed and that seemed involved in cytopharinx and oral mouth neogenesis. The investigation was focused on the V-ATPases generating H^+^-gradient across membranes. The V-ATPases are present in abundance in the CV membranes of different protozoa living in a wide range of osmotic environments and important for vacuolar function in expelling water entering the cells by osmosis (Allen and Naitoh; 2002). We first checked if V-ATPases were localized at the CV. We proceeded by immunostaining of cysts and teletroch with a polyclonal antibody against the B1-subunit of the V1-ATPase subcomplex. The results established that in *Vorticella* as in other protists these proteins are plasma and vacuolar membrane-resident (Fig. 7A-D). The typical cressent-like staining of a single vacuole among the food vacuoles and the contractile one strongly indicated that the antibody decorated membranes of the CV. Next, we examined the action of the concanamycin A, a specific inhibitor of the V-ATPases, on cyst differentiation and release. Cysts in stationary phase were collected, washed and then incubated in water in the presence or not of 5 µM concanamycin A. We therefore examined the action of the drug on the function of the CV complex in cyst differentiation and release based on counting in each condition both (i) filled and versus empty cyst walls to calculate the ratio between empty and total cysts numbers (Fig. 7E); and (ii) mean speed of telotrophs in motion (Figure 7F). As expected, no empty cyst wall was detected in cysts at stationary phase (SP; 14-day culture in V8 medium) while a high rate for empty cyst walls was calculated in the control (C) after one hour of release in water. In contrast and at the same time point, in concanamycin A-treated samples (Conca A) the rate of empty cysts was drastically reduced compared to control cells (Fig. 7E). Concomitantly, there was also a marked decrease in the mean speed of moving cells in the presence of the drug (Fig. 7F) indicating that cysts were impaired in the differentiation and/or release processes. Next, we visualized α-tubulin by immunostaining. In the control cells, the α-tubulins were mainly distributed at the two poles of most released telotrochs (Fig. 7G) drawing the characteristics two ciliary arrays of *Vorticella* sp. which promote fluid flow for cell motility; In the concanamycin A-treated cells, cysts and telotrochs were faintly stained compared to the control cells (Supplementary Fig. 3 A,B). The telotrochs could exhibited a staining more uniformly distributed and not reminiscent to a clear polarized status indicating that ciliogenesis, involving polarized basal body docking onto cell membranes, were inhibited or slowed (Fig. 7G). A larger CV could also be observed with an extended systole and no structure reminiscent to the cytopharingal cavity nor ciliated oral mouth (Supplementary Fig. 3, H-L).

**Figure 7.**
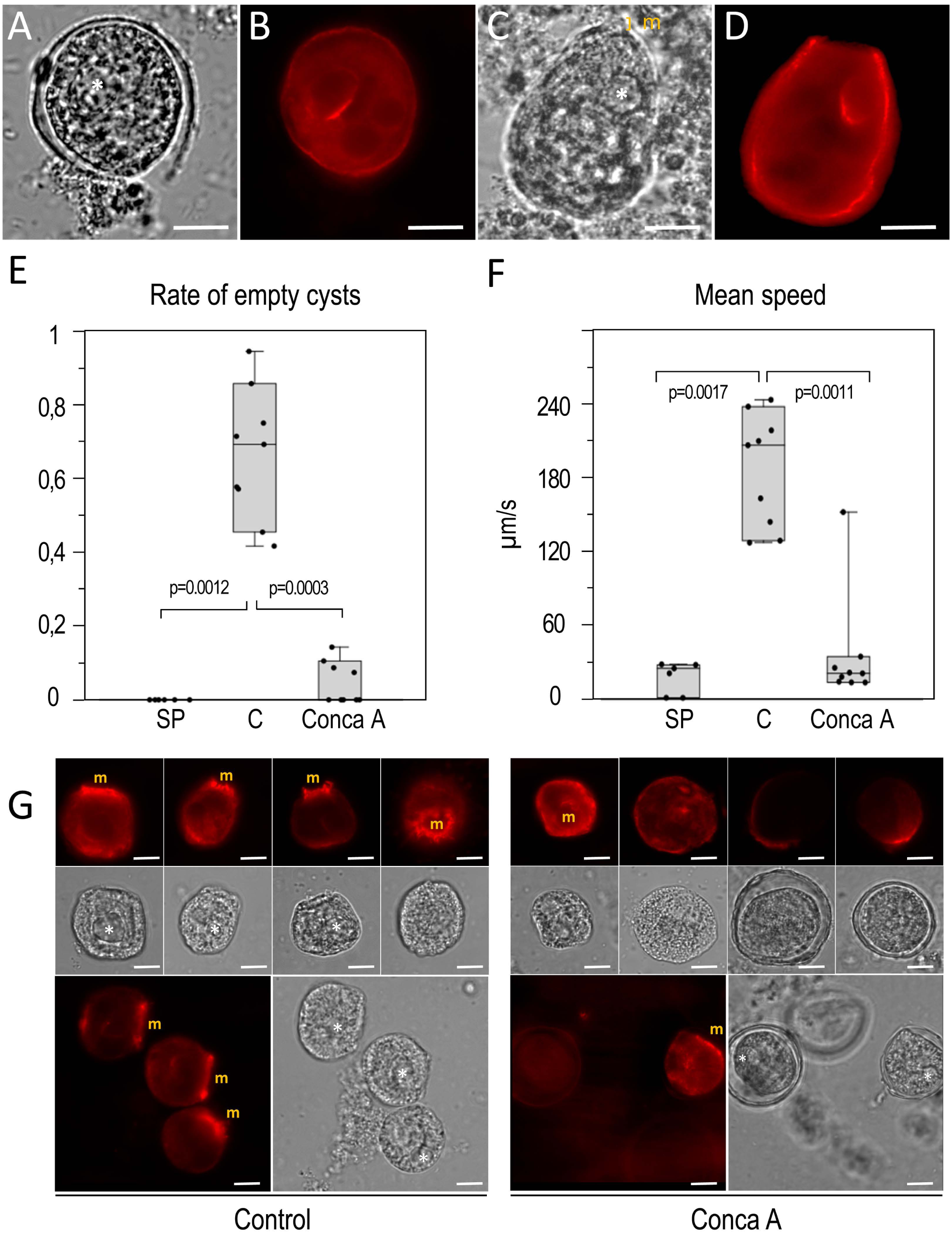
Concanamycin inhibits differentiation and release. (**A-D**) Phase contrast (**A, C**) and immunostaining (**B, D**) of a cyst and a teletroch with a polyclonal antibody against the B1-subunit of the V1-ATPase subcomplex. (**E**) Rate of empty cysts and (**F**) telotroch mean speed, at stationary phase (SP), after one hour of release in water in the presence of concanamycin A (Conca A) or not (C for control). (**G**) Phase contrast and α-tubulin immunostaining of representative cysts and teletrochs treated or not with concanamycin A. White stars and “m” annotation indicate the CV and oral mouth position, respectively. Scale bars, 10 μm.

## DISCUSSION

There is today a real Vorticella paradox. Although it was probably the first microorganism described in detail (Van Leeuwenhoek, 1677), *Vorticella sp.* remains a “poor little creature” being among the most neglected organisms in terms of available data in cellular and molecular Biology and Genomics. Only 328 nucleic acid and 9 protein sequences are accessible in databases such as NCBI or Uniprot. It seems not to have yet appeared in the Protist 10 000 Genomes Project (Gao *et al*., 2024). And yet, its distinctive single-celled forms (cyst, telotroph, trophont) make them really suitable objects of investigation on the mechanisms that govern biological transitions within a species. Using microscopy and pharmacology, the current study explored the cyst-telotroph transition in the ciliate Vorticella allowing to excystment.

### The cyst excystment of the Vorticella ciliate

(Li *et al*., 2022) have reviewed a vast amount of information available on changes and some known mechanisms involved during excystment process of 14 representative ciliated protists. General principles on which excystment is induced involved first reappearance of ciliary and intracellular structures that had degenerated during encystment and before excysting. Second, escape from the cyst wall is assisted by a preformed or neoformed pore on the cyst wall and a rupture generated by the pressure of cell movement inside the cyst wall and excystation vacuole. In *Pleurotricha lanceolate,* generation of the pore occurred at uncertain position (Jeffries, 1956).

In the current study the data indicate that the papilla and the CV orchestrate a complete differentiation in telotroch from a cyst initially constituted of unstructured hyaloplasm/cytoplasm. They allow us to define two patterns characterizing the cyst transition to telotroph. A first temporal pattern is initially marked by the functional formation of the CV, which then discharges its fluid into the neoformed cytopharynx and, in a third stage, into a membrane invagination which, to the rhythm of the vacuole, differentiates into an oral cavity. The second pattern, a spatial one, is delineated on one pole by the position of the papilla, which determines the site of oral cavity formation and ciliogenesis, directly below it and dictated by the CV tempo. The point of cyst wall rupture, leading to the release of the telotroch in an apical-basal polarity defines the second pole and is characterized by β-actin accumulation. The data emphasizes the role of CV and papilla in the two axes setting up (Fig. 8).

**Figure 8.**
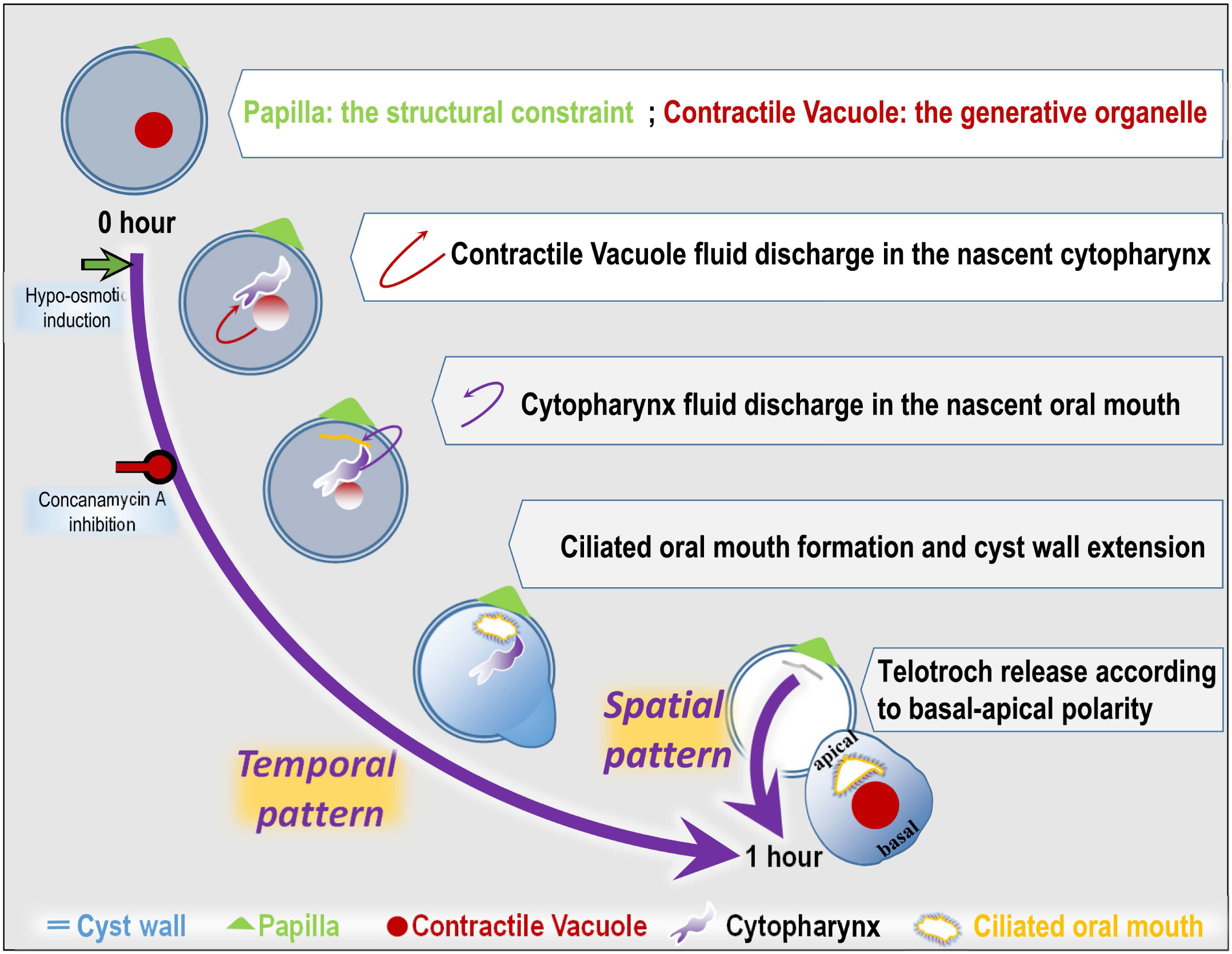
**Overview of events that contribute to the formation of telotroch release from a cyst in *Vorticella microstoma*** A step-by-step schematic diagram of a cyst/telorotroch transition depicting the hypothesis raised in this work according to which within the cyst, the papilla position at the wall cyst and a functional contractile vacuole cause and orient organelles formation (cytopharynx and ciliated oral mouth) as well as apical-basal telotroch polarity, and subsequent telotroch release.

Indeed, two lines of evidences designate the CV if not the conductor of the establishment of the first temporal pattern, at least as a key upstream player. (i) It is the first functional organelle to emerge. At the strict tempo of its contractions, the appearance of the cytopharynx and then the oral mouth sequentially follow. (ii) Concanamycin A treatment compromising its V-ATPase-dependent functions alters ciliogenesis, polarized ciliary arrangement, release and motion of telotrochs. CV could act by regulating osmotic and ion homeostasis as suggested by the increase in size of released telotroph compared to cyst. It could also act by discharging signaling molecules to the plasma membrane of nascent organelles (cytopharynx, oral mouth), *as D. discoideum* CV discharges cAMP and Ca^2+^ to plasma membrane at the rear of each cell during early cell aggregation allowing stalk formation (Fadil and Janetopoulos, 2021).

A passive but key player in the spatial pattern definition is clearly the papilla. Its position initially defines the plasma membrane anchored one where oral mouth will emerge precisely positioned neatly under the papilla. By this way it determines the position of the apical-basal polarity of the neoformed telotroph. Considering that the release always occurred according to the basal-apical polarity of telotroph and within the semi-circle opposite to the papilla, papillae must be considered as the organisers establishing the spatial pattern of excystment. Thus, a papilla position is a reference point of a developmental process and not a simple morphological ornament precious for taxonomy.

### Morphogenesis of apical-basal polarity

More broadly, telotroch differentiation in Vorticella should also be a new model to investigate the apical-basal polarity that is essential to many morphogenetic processes in eukaryotes. Its establishment is extensively studied at the genetic and molecular level in higher eukaryotes (Motegi *et al*., 2020), in ciliates (Soh and Pearson, 2022) or amoebozoa as the unicellular *D. discoideum* (Fadil and Janetopoulos, 2021). In *Vorticella* the establishment of apical-basal polarity is achieved within the cyst before release and simply results of a single-cell process primary involving the same two structures, the generative CV and the positional papilla. The pre-existing papilla of the cyst wall is a structural constraint defining the position of the apical organization of future telotrophs through attachment of a particular cortical area of the plasma membrane from which emerges the oral cavity. This could be view as an additional facet of “structural inheritance” whereby in ciliates pre-existing structures enforce a developmental plan for newly formed structures (Beisson & Sonneborn, 1965). In *Tetrahymena pyriformis* and *Paramecium tetraurelia*, the pre-existing intracellular cortical organization of cilia deliver information for the formation and positioning of new structures (basal bodies, microtubular network) defining the cortical organization after scission (Beisson & Sonneborn, 1965; Ng and Frankel, 1977; Soh and Pearson, 2022). It is also reminiscent to maternal tissues acting in higher eukaryotes as a mechanical constraint (Woudenberg *et al*., 2024) as the cell wall-mediated maternal control of apical–basal patterning in brown algae (Shaw and Quatrano, 1996; Dries *et al*., 2024). The present data clearly indicate that in *Vorticella*, the papilla constitutes the extracellular anchorage point for cellular material assembled during encystment and pre-required for oral cavity and cilia array formation during excystment. Such material could include the fibrillary one viewed after release in Fig. 2F and Movie 1, still linked at the papilla site after release, and conforming to the shape of the thin sheet forming beneath the papilla (Fig2C). The establishment of apical-basal polarity also involves a functional generative CV complex. The analyses of the movies clearly establish that CV expels its fluid through the pore in the cytopharynx. The discharge is then mainly propagated until the papilla-located thin sheet contributing to oral cavity and cilia array formation. The discharge wave did not propagate to the rest of the cell which constitute schematically below the vacuole, the basal side of the cell. Thus cellular process that is happening, it is as though fluid discharge defines a developmental apical way from the CV to the papilla on which cytpharynx and oral mouth genesis take place.

## Conclusion

Encystment and excystment are fundamental to the conservation and dissemination of protists in the biosphere. Most of the molecular and cellular associated mechanisms remains elusive. We highlighted generative and structural elements that organise from a cyst and according to a spatio-temporal plan the telotroch differentiation, apical-basal cell polarity and excystment in the ciliate *Vorticella*. By the identification of the papilla as a positional constraint and the CV as a generative organelle of cell polarity in a single-cell system, we provide a new and additional evidence that apical-basal cell polarity is an ancestral trait not only widespread among eukaryotic multicellular cell organisations. Future studies in cell biology will be required to understand how the papilla is formed during encystment, which cellular components are locally recruited to precisely mounted the site of oral cavity and apical cilia array formation, and which CV function(s) (water exclusion, ion homeostasis, signaling molecule driver) controls the cell differentiation.

## MATERIALS AND METHODS

### Vorticella Strain and sample treatment

The *Vorticella microstoma* 30897 stain, in co-culture with *Enterobacter aerogenes (*13048 strain), was purchased from the American Type Culture Collection (ATCC) collection of protists (LGC standards). The cells were cultivated for 3 to 14 days in V8 liquid medium at 24°C. Three days were necessary to observe a high rate of swimming telotroch. Ten to fourteen days were required to reach a 100% rate for cyst formation. In the intermediate range of time, the trophont appeared to be the most common form observed. To induce cyst excystment by means of a hypo-osmotic stress, the medium of 10-14-day-old cultures was discarded and cysts were resuspended in water (the time point t=0 in this study). Samples (100-200 µl) were rapidly mounted in a chamber made of a microscope slides and a top coverslip loaded on two spacers (0,5-1 mm) for starting microscopic observation soon as possible.

To inhibit the V-ATPase, cysts were treated with Concanamycin A, a specific inhibitor of V-ATPases binding to the proteolipid subunit c of the V0 complex (Huss *et al*., 2002). To assess the function of the CV machinery we proceeded as previously described by Tesmesvari *et al*. (1996) with *Dictyostelium discoideum*. Cysts in the stationary phase were harvested by centrifugation, resuspended in water to create a hypo-osmotic environment and treated or not with 5µM Concanamycin A in solution in DMSO. The appropriate DMSO dilution was applied to the control samples.

### Microscopy

Images (size 512×512 or 1029×1044 pixels) and movies (image 512×512 pixels; time scale 0.2 or 0,4s; time series 1000-3000 s) were obtained using a Zeiss Axio Z1 microscope equipped as followed : a plan-apochromat63x/1.40oil M27 and a EC-plan-neofluofluar 40x.75 M27 objectives; a Hamamatsu camera and the ZEN 2012 pro lite application (Carl Zeiss Microscopy, Germany). The material was available on the Microscopy Platform-Sophia Agrobiotech Institut-INRAE 1355-UNS-CNRS 7254-INRAE PACA, Sophia Antipolis.

### Immunohistochemistry

Cysts and telotrochs were collected by centrifugation. They were fixed in methanol for 10 min at −20°C and then incubated in phosphate-buffered saline (PBS), pH 7.2 at room temperature. Samples were treated in PBS with 0,1% Triton for 20 min at room temperature, and later with a blocking solution containing 3% of bovine serum albumin in PBS, for 30 min. They were then first incubated for 18h at 4°C with a rabbit polyclonal antibody against the α-tubulin subunit (NeoBiotech), or against the V1-ATPase B1 subunit (NeoBiotech), or against the β-actin diluted 1:200. Samples were washed three times with PBS and incubated for 1 h at room temperature with a FluoProbes 547H Donkey anti-Rabbit IgG or a Fluoprobes 488 anti-rabbit IgG antibody diluted 1:200 (Interbiotech – BioScience Innovations). Samples were rinsed three times and mounted in Fluoroshield with DAPI (Sigma, St. Louis, MA USA).

### Image time series analysis

To analyse CV beating, cytopharynx and oral mouth expansion we proceeded by kymography using in FIJI (Schindelin *et al*., 2012) the Measurement_Tool plugin (k**-**button; https://dev.mri.cnrs.fr/projects/imagej-macros/wiki/Velocity_Measurement_Tool). To generate the kymograms, we applied the tool by selecting segmented lines including a diameter and the centre of the vacuole or delimiting as much as possible the mid-section of the papilla, oral cavity, cytopharynx and contractile vacuole. In the kymograms, space was measured along the x-axis while time was measured along the y-axis. To determine mean velocity of cysts and telotrochs, cells were tracked with FIJI using the TrackMate plugin (Ershov *et al*., 2022) as per the procedure detailed in Lupatelli *et al*. (2023).

### Statistics

Experiments were repeated at least 3 times. Data are presented as mean values±SEM. One-way ANOVA with Mann-Whitney pairwise test, were used for the statistical analysis. Past4.17 software was used to analyse the data. Bonferroni corrected/adjusted p values of <0.05 are indicated and were considered.

## Author contributions

EG and MLK conducted the analyses. EG wrote the first draft. AA and MLK contributed to the writing of the manuscript. EG and AA managed acquisition of the financial support for the project leading to this publication.

## Acknowledgements

A warm thank-you goes to Carlotta Aurora Lupatelli for very fruitful discussions on the topic of the paper.

---

**Supplementary Figure 1**

**Telotroch Release from cyst**

Micrographs of six cysts and telotrochs at t=0 (left) and approximately t=50 min (right) just after release. (p) indicates the papilla localisation, and (*) the position where the cyst wall will be (left) and is ruptured (right). The apical-basal polarity of telotrophs is indicated immediately after release illustrating that excystment involves a specific orientation of telotrophs in the opening of the cyst wall with the basal end at first and then the apical end (easily identified by the oral cavity). Scale bar: 20 µm.

**Supplementary Figure 2**

**Kinetics of vacuole beat frequency as a function of the ordered series of diastole-systole cycles**

Nine independent time-lapse experiments recording cysts from t=0 to the release time ranging between 20 and 40 min. X-axis indicates the serial number of cycles; y-axis is indicative of the beating frequency between two successive pulses.

**Supplementary Figure 3**

**α-tubulin staining of cells treated or not treated with concanamycin A**

(**A-B**) Merged images of phase contrast and α-tubulin immunostaining of representative cysts and teletrochs treated (**B**) or not (**A**) with concanamycin A. Scale bar: 50 µm. (**C-G**) typical timing of systole duration of 1s with the dynamics of engulfment of the cytopharynx and emergence of oral cavity signed by membrane invagination below the papilla. (**H-L**) Extension of systole duration (7s) with no structures reminiscent to the cytopharingal cavity nor cilated oral mouth in a concanamycin A-treated cyst connection. Scale bar: 10 µm.

**Movie1 - Time-lapse video microscopy of a cyst differentiating in a telotroch**

Images were taken every 500 ms with a 40× objective. t=0 was undetermined. Annotation recordings is frame-by-frame.

**Movie2 - Video microscopy of a cyst differentiating in a telotroch**

Time-lapse imaging with 0.4-s intervals. Images were taken every 400 ms with a 40× objective. The indicated timing is refered to t=0 when the differentiation started in response to hypo-osmotic conditions.

**Movie3 - Video microscopy of a cyst differentiating in a telotroch**

Images were taken every 200 ms with a 40× objective. The indicated timing is refered to t=0 when the differentiation started in response to hypo-osmotic conditions.

