## Supplementary figures and images for "Contractile Vacuole and Papilla drive Cyst/Telotroch transition in *Vorticella microstoma*"

### Supplementary Figures 1-3

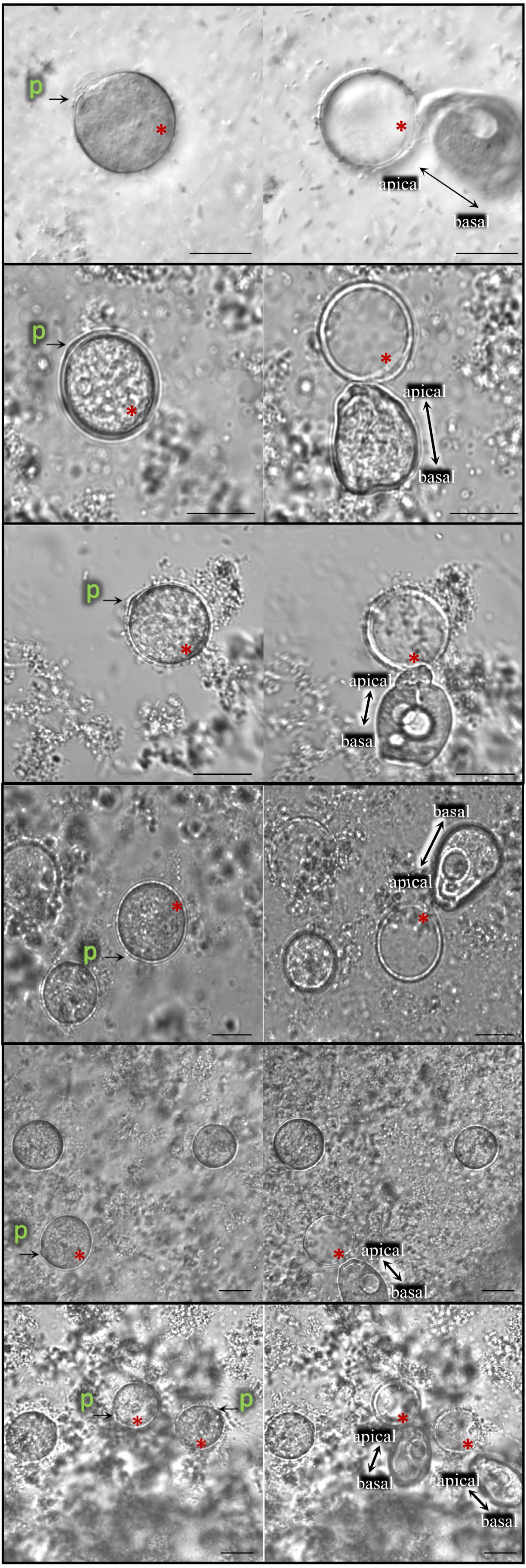


#### Supplementary Fig 1

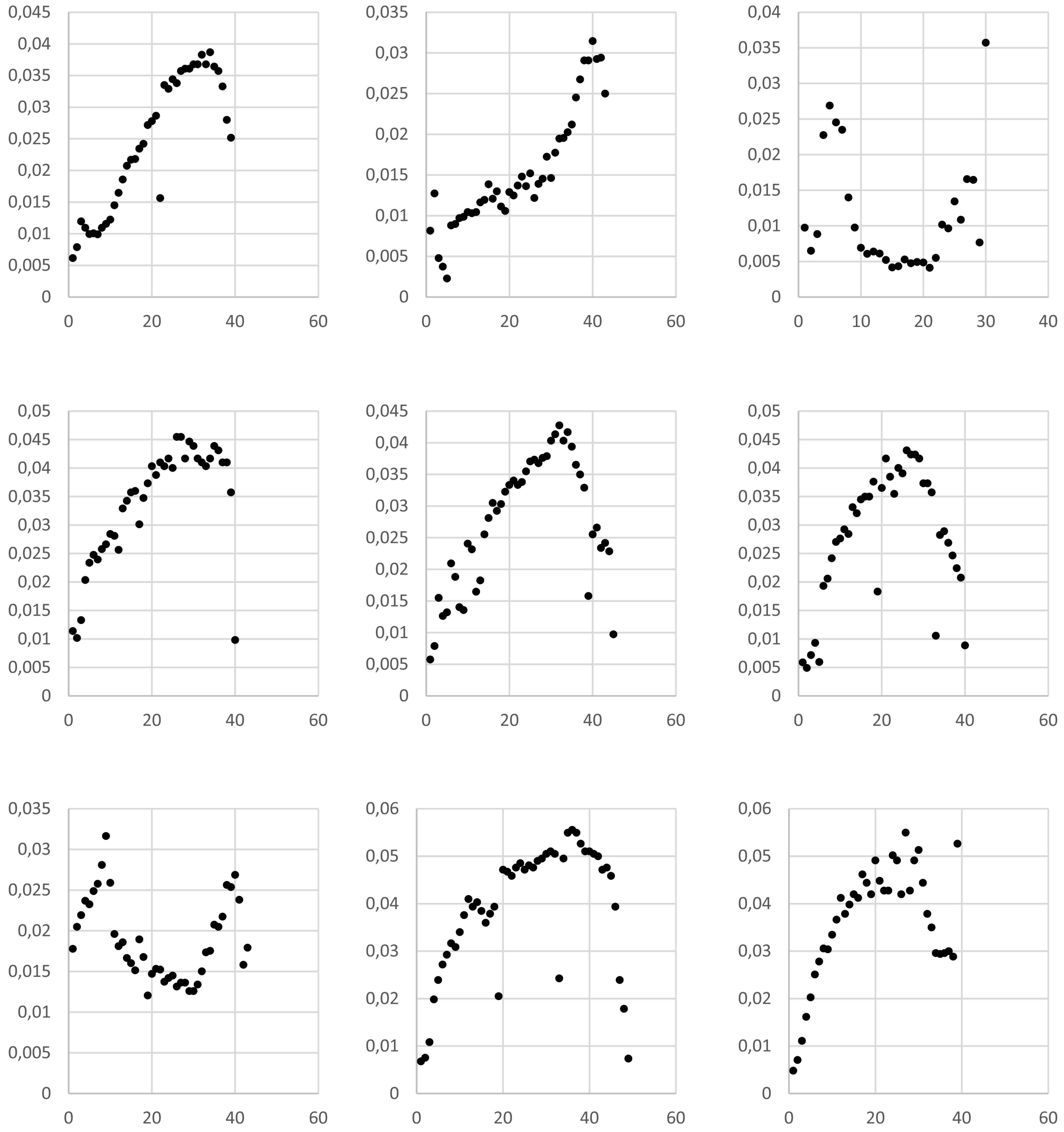


#### Supplementary Fig 2

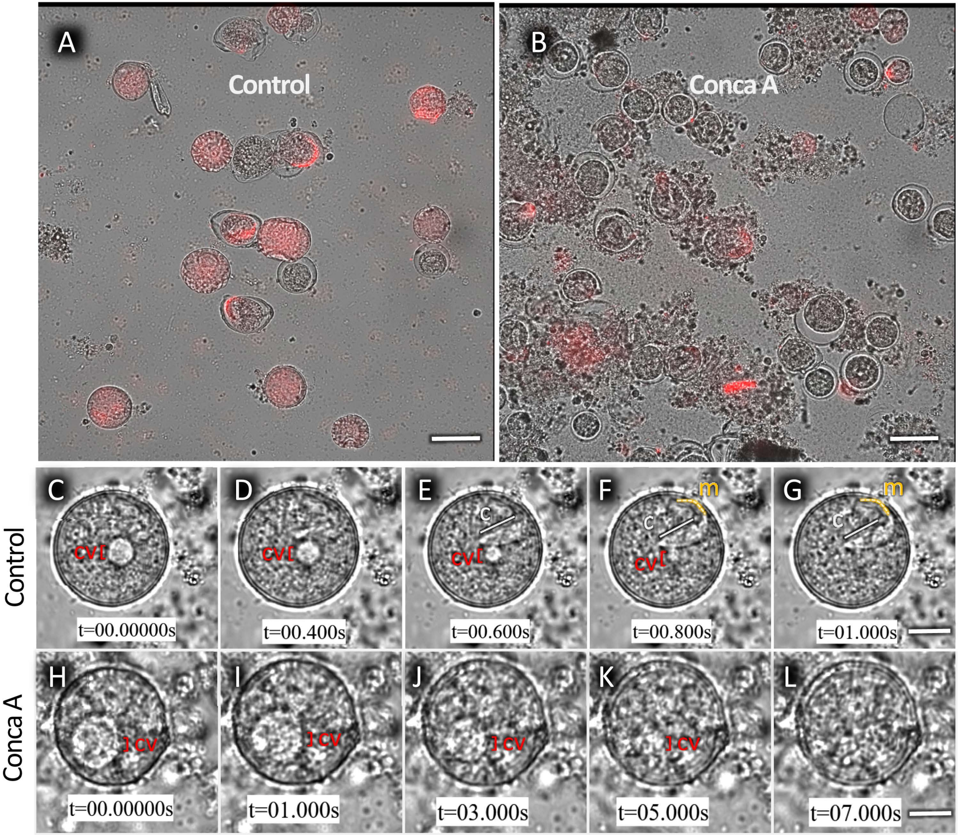


#### Supplementary Fig 3
